# MP-NeRF: A Massively Parallel Method for Accelerating Protein Structure Reconstruction from Internal Coordinates

**DOI:** 10.1101/2021.06.08.446214

**Authors:** Eric Alcaide, Stella Biderman, Amalio Telenti, M. Cyrus Maher

## Abstract

The conversion of proteins between internal and cartesian coordinates is a limiting step in many pipelines, such as molecular dynamics simulations and machine learning models. This conversion is typically carried out by sequential or parallel applications of the Natural extension of Reference Frame (NeRF) algorithm. This work proposes a massively parallel NeRF implementation which, depending on the polymer length, achieves speedups between 400-1200x over the previous state-of-the-art. It accomplishes this by dividing the conversion into three main phases: parallel composition of the monomer backbone, assembly of backbone subunits, and parallel elongation of sidechains; and by batching these computations into a minimal number of efficient matrix operations. Special emphasis is placed on reusability and ease of use. We open source the code (available at https://github.com/EleutherAI/mp_nerf) and provide a corresponding python package.

## Introduction

Molecular modelling often employs two distinct sets of coordinates to represent proteins or other polymers, such as nucleic acids or glycans. One can represent a polymer using internal coordinates (bond lengths, bond angles and dihedrals) or cartesian coordinates (x,y,z)^1^. Either of these two coordinate systems allows for a complete representation of the molecule, and both have their strengths and weaknesses. Cartesian coordinates are easier to work with when treating the polymer as a rigid system (e.g. for rotations, translations or visualizations), whereas internal coordinates are preferred when working with forces and interactions between the polymer atoms.

Due to this complementarity, it is often necessary to convert between the two representations. Translation between these systems, however, can become a bottleneck in many applications. To align with prior work, we will refer to the transform from cartesian to internal coordinates as *forward translation*^2^. On modern hardware, this process can be straightforwardly parallelized across all polymer points, since the internal coordinates can be calculated independently for each point. This is not the case for *reverse translation,* however. Translation from internal coordinates to cartesian ones has typically been carried out sequentially, since the position of each atom depends on the position of the previous one^3^.

This sequential dependency bottlenecks reverse translation. Although the calculation can be parallelized across many polymers of similar length, this bottleneck is significant for applications that make intensive use of forward and reverse translation. Examples of such applications include the training of machine learning models^4^, protein structure refinement from NMR data^1^, analysis of protein structure changes^3^, and molecular dynamics simulations^5^.

With the development of more and better computational tools, some effort has been devoted in recent years to alleviating the reverse translation bottleneck^4,5^. These works have focused on the usage of high-performance, optimized code; the division of the backbone into different fragments (which are folded independently and later ensembled)^4^; and tree ensembling algorithms^5^, among other strategies. These approaches, however, are often implemented specifically for Graphical Processing Units (GPUs), thus limiting usage to expensive and specialized hardware accelerators. Despite these improvements, the translation from internal coordinates to cartesian ones continues to be a bottleneck even in the many GPU-based pipelines.

The standard algorithm used for reverse translation is the Natural extension of Reference Frame (NeRF) procedure^3^. The pNeRF algorithm^4^ explored extending NeRF by dividing the polymer into different fragments, iteratively folding for each fragment in parallel, and then concatenating the independently folded fragments. However, it considered only the backbone, and the number of parallelized fragments was low (on the order of 1-10). A more recent implementation of a parallel NeRF algorithm^5^ considered the sidechains as well, and performed a tree-based merge of the different fragments. However, it required specialized hardware such as CUDA-capable GPUs and there is friction when trying to adapt the implementation for different usecases.

This work builds on top of the theoretical proof of correctness shown in previous work^3^, and also provides numerical validation through error analysis in back-and-forth translation cycles as in^4^. We introduce a massively-parallel (mp-NeRF) algorithm, which reorders the steps in the parallel implementations of the NeRF algorithm to unlock parallelization and optimize execution when possible. The algorithm leverages ideas from previous work^4,5^, and builds on top of them to provide an acceleration of over 400x-1200x, depending on the length of the protein being translated. It accomplishes this by decomposing operations into parallelizable units and using more efficient data structures that leverage the efficiency of matrix-based operations. This algorithm thereby increases the throughput of pipelines which make heavy use of such conversions, such as the training of machine learning models to predict protein folding.

## Methods

### Massively parallelized natural extension of reference frame

Many polymers, such as large biomolecules, can be divided into a backbone and a set of offshoot branches, usually referred to as sidechains. This work initially takes a parallel scheme similar to pNeRF, which only considers the protein backbone and a small number of fragments to build in parallel, but takes it to the extreme by considering the backbone of each amino acid as a separate fragment. It also extends pNeRF by incorporating side chains. Since the polymer backbone is composed of repeated subunits, it is convenient to parallelize computation across monomer backbone fragments. The procedure can be separated into the 3 phases described in detail below. Briefly, these encompass folding backbone subunits, linking these subunits, and then folding side chains.

#### 1. Parallel composition of the minimal repeated structure

For every polymer backbone subunit, we initialize two points near the origin coordinates. Together with the origin point, these three points define a plane that can be used to define the first dihedral angle and initialize polymer extension. In subsequent steps, this reference frame is also used to calculate the relative orientations needed to connect monomer backbones properly. From there, we implement the NeRF algorithm sequentially for every atom in a given monomer backbone, until we reach the first point of the next monomer backbone, as illustrated in **Fig. 1A**. This process is repeated in parallel for all monomers. This step requires a number of NeRF calls equal to the number of atoms in the backbone monomer, times the number of structures across which the calculation is parallelized. A theoretical proof of the equivalence of this step with respect to sequential elongation can be found in AlQuraishi 2019^4^.

**FIGURE 1:**
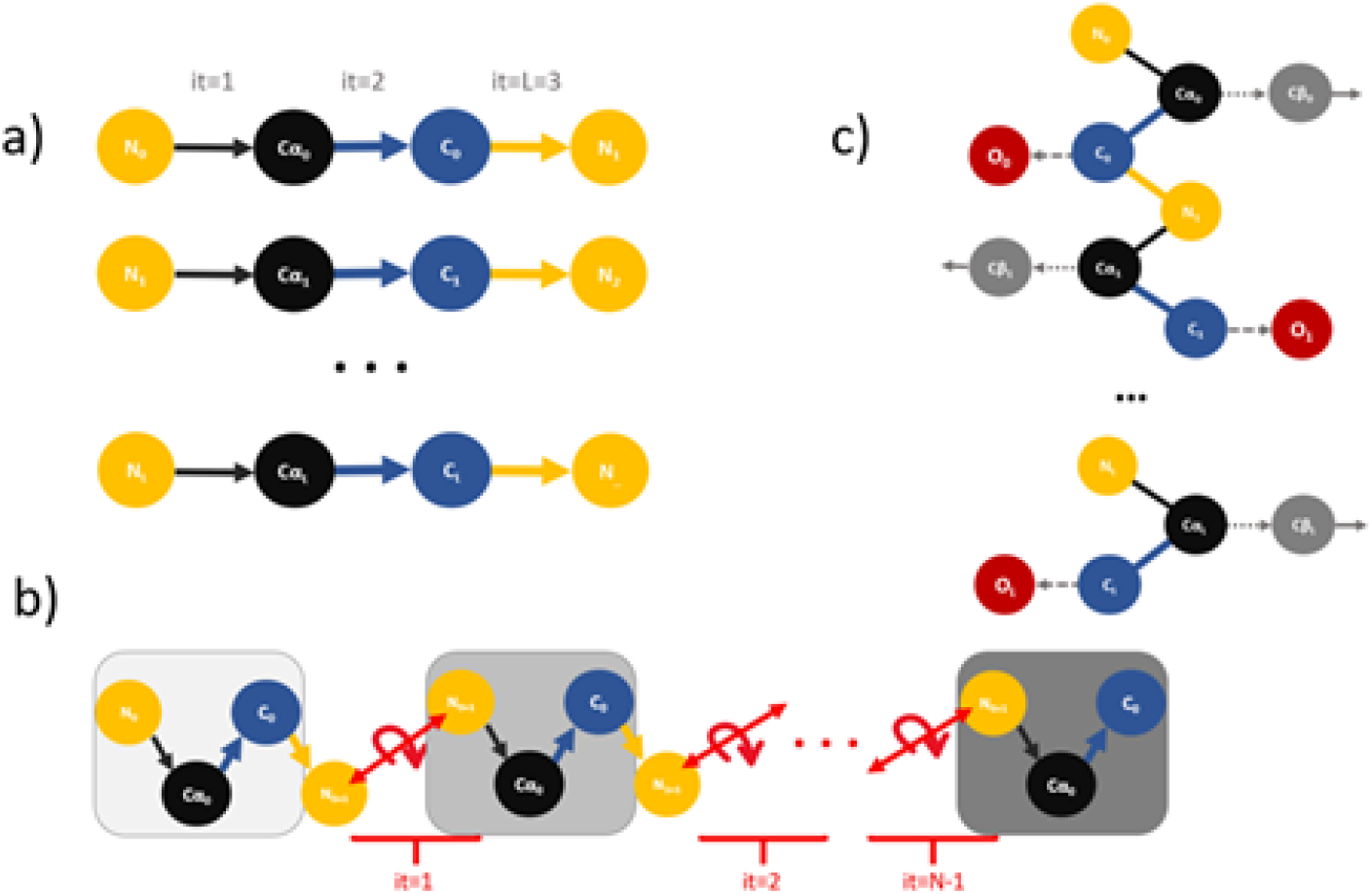
Illustration of the algorithm. a) Parallel composition of the monomers. b) sequential concatenation of monomers. c) Parallel composition of side chains.

#### 2. Assembly of backbone monomers

We then join the backbone monomers by an efficient roto-translation operation. That is, we move the points of each subunit so that its first atom is now linked to the last atom of the previous monomer, and we rotate the assembled subunits so that their orientation matches the reference frame of the structure. This sequential pass requires:

1. Batched matrix multiplications to construct the rotation matrices, equal to the number of subunits to assemble
2. An iterative matrix multiplication to rotate all subunits sequentially
3. A cumulative sum of length *N* − 1 given by the expression 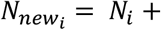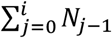 where *N_j − 1_* represents the last point of the *j*th subunit after rotation and *N_new_* represents the roto-translated backbone subunits. This process is illustrated in Fig. 1B.

#### 3. Parallel composition of side chains

After the backbone assembly, we perform the calculation of the side chains in parallel, as illustrated in **Fig. 1C.**This requires a maximum number of NeRF calls equal to the maximum number of atoms in any possible side chain, times the number of side chains across which the calculation is parallelized. Note that this is an upper bound, since not all sidechains will have the same number of points.

### Implementation details

The calculations for the rotation matrices to join the backbone fragments are decomposed into a rotation from the reference frame to the first monomer backbone (*M*_0_), and from each monomer backbone to the next one in the chain (*M*_i_). This allows for the parallel construction of rotation matrices, leaving a cumulative matrix multiplication and a cumulative sum as the only sequential parts of the algorithm:

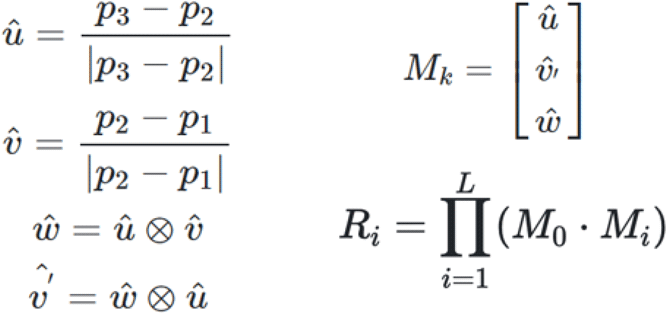

When constructing the *M_i_* matrix; *p*_1,_ *p*_2,_ *p*_3_ are, respectively, the coordinates of the carbonyl and alpha carbon atoms of the present amino acid, and the nitrogen atom of the next amino acid. When constructing the *M_0_* matrix; *p*_1,_ *p*_2,_ *p*_3_ are the coordinates for the two points adjacent to the origin, and the origin itself. These define the reference frame. In the expression above, *M_k_* represents both the *M_i_* and *M_0_* cases.

The translation operation is implemented as a cumulative sum, accelerating its calculation. Special effort is put into generalizing every possible function for an arbitrary number of atoms, so that a single function call can do the required calculations for as many atoms as possible, thus achieving a near-perfect usage of the processor native parallel capabilities (CPU-native vector instructions such as SIMD and AVX or GPU massively parallel architecture).

Protein-specific optimizations include omitting the oxygen atom in the backbone carbonyl, since it can only have one orientation relative to the carbon to which it is bonded. We later incorporate it as a sidechain addition to the main backbone formed by the N-CA-C atoms of each amino acid. During the translation process, the rings present in amino acid sidechains are simplified so that there is a unique path connecting every atom in the sidechain to the backbone. We do this by masking one bond per ring, so that only linear and branched sidechains are considered, and thus there are no loops in the algorithm’s reconstruction path. This masking does not require further processing in the output, since the algorithm returns a point cloud, consistent with PDB format. For data processing, the *sidechainnet^6^* format is used to encode proteins as arrays of shape *(N × 14 × 3),* where N is the length of the protein.

A profiling report is included in the Supplementary Information section (**Figure S1**).

## Results

To emphasize the power and efficiency of our methodology, we compare to both the state-of-the-art4 (SOTA) and the “state-of-practice,” i.e., the most advanced techniques that are currently in widespread use. A presentation of the compared algorithms and details about their implementations can be found in **Table 1**.

**Table 1.**
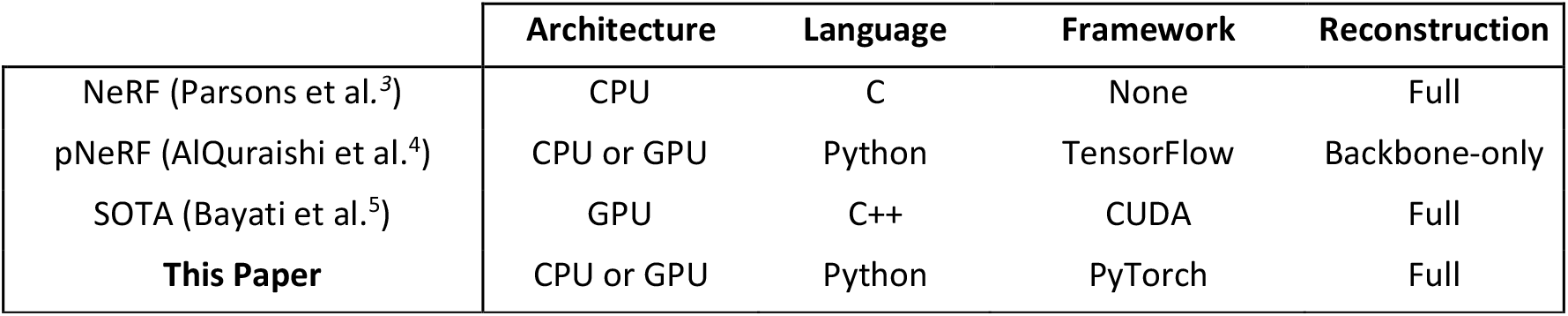
Different algorithmic implementations and hardware requirements.

|  | Architecture | Language | Framework | Reconstruction |
| --- | --- | --- | --- | --- |
| NeRF (Parsons et al.<sup>3</sup>) | CPU | C | None | Full |
| pNeRF (AlQuraishi et al.<sup>4</sup>) | CPU or GPU | Python | TensorFlow | Backbone-only |
| SOTA (Bayati et al.<sup>5</sup>) | GPU | C++ | CUDA | Full |
| **This Paper** | CPU or GPU | Python | PyTorch | Full |

### Computational Efficiency

Our algorithm achieves more than 400x-1200x improvements over the previous SOTA for proteins, depending on the protein length studied. It does this using only CPUs, which makes it broadly applicable and able to exploit parallelism in CPU cluster setups. It can be seen in **Table 2** that the GPU implementation is slower than its CPU counterpart for all values of polymer length. This might be due to the cost of data transfers and synchronization between CPU and GPU, the cost of spawning GPU kernels for each step, the lower clock frequency of the GPU, which notably slowed down the sequential assembly of backbone monomers, and an under-utilization of its parallelism. Although proteins longer than 1000-AA are infrequent, the GPU implementation might show advantage in other, longer polymers such as nucleic acids. As evidenced in **Table 2**, this massively parallel algorithm eliminates the need for hardware accelerators for this task, lowering the requirements to perform efficient conversion of polymers from internal to cartesian coordinates.

**TABLE 2:**
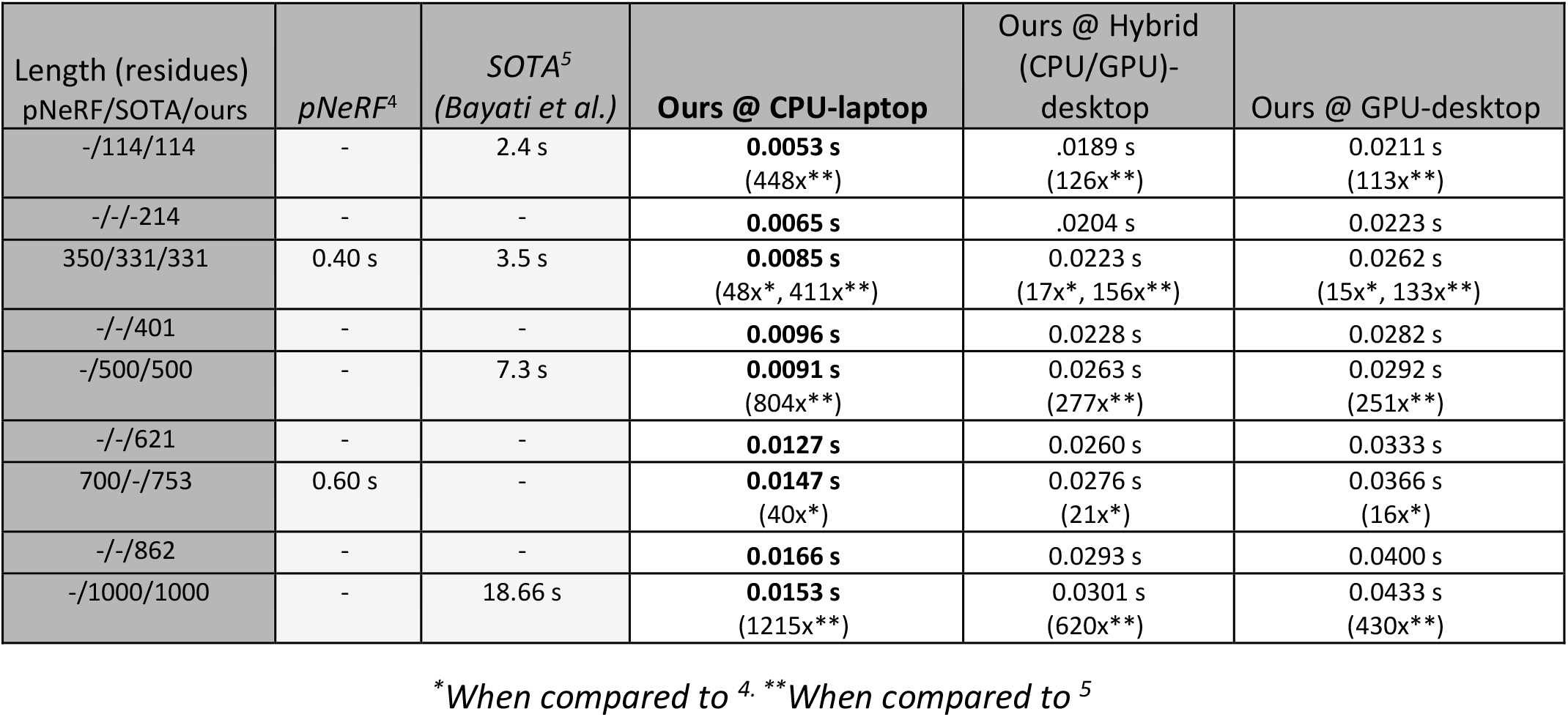
Comparison of execution times between previous SOTA, pNeRF, and our implementation. We report results from the GPU version of pNeRF. For our implementations, speedups are reported between parenthesis and with the “*– x”* suffix expressing speedup with respect to pNeRF and the recent SOTA implementation. Note that the pNeRF implementation only considers the 3 backbone atoms N, C_α_, C, whereas the other algorithms consider the full side chain. Due to the difficulty of benchmarking all algorithms on the same proteins, we conservatively compare to comparably sized, but longer proteins.

### Accuracy

We check the cumulative error that comes with repeated transformations from internal to cartesian and back again. This round-trip is of special importance in algorithms that use internal coordinates, but that need conversion functions before and after since the molecular simulation or base representation is in cartesian coordinates. A variation in the error during the first 100 out of 1000 forward-backward cycles of conversion can be observed. We hypothesize this might be due to numerical inaccuracies in floating-point arithmetic. Importantly, we note that errors do not significantly accumulate even over 1000 roundtrip translation cycles.

**FIGURE 3:**
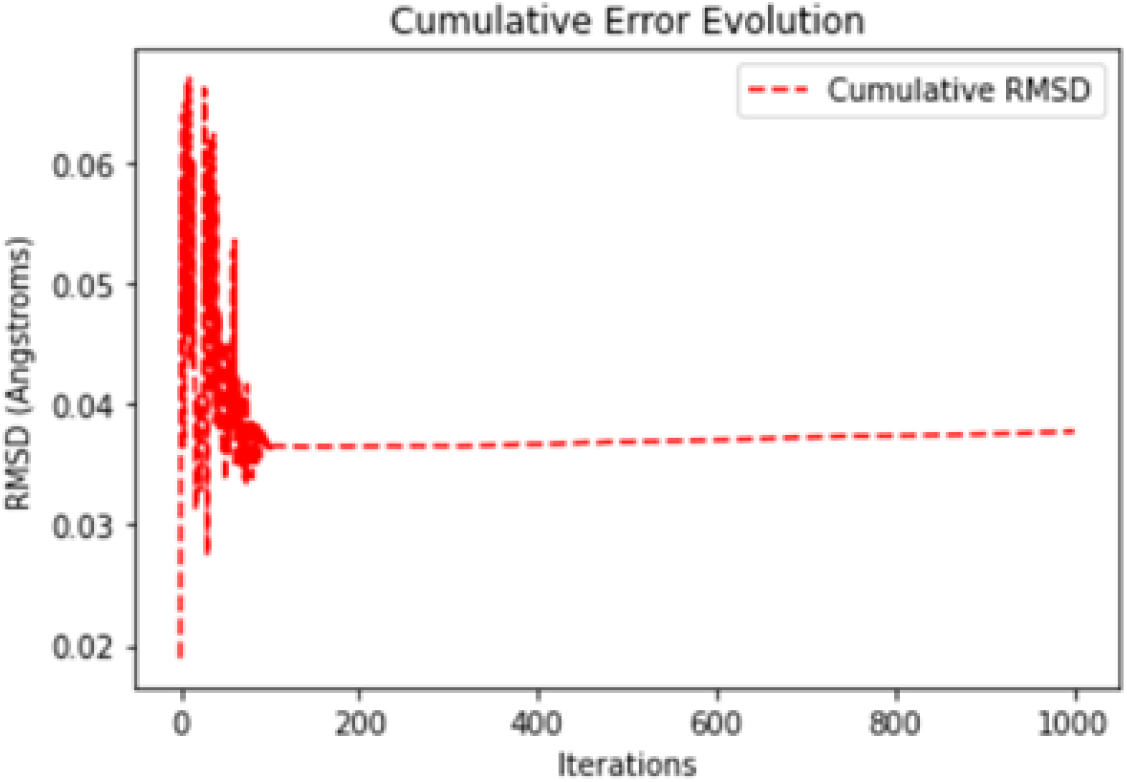
Cumulative mean error (RMSD) as a function of encoding-decoding cycles.

### Design Tradeoffs

We would like to emphasize that some design choices are currently limiting the speed of the implementation presented here. However, we accept these tradeoffs to ease use and facilitate adoption. These design choices include implementing the algorithm in Python instead of a compiled language. Python is a high-level language widely known among the scientific community, with many scientific software packages implemented in it. Since Python is an interpreted language, functions that are not ultimately implemented in more performant languages like C++ can be slow. However, since the majority of our execution time is devoted to matrix operations implemented in C (Figure S1, Table S1), we do not incur a large performance penalty in Python. We estimate that the current implementation could be accelerated by 2x if we switched to a compiled language. However, this would inevitably lead to a reduction in adoption, increased cost of code maintainability and a reduction on the possible extensions, adaptations, reusability, and readability of the current implementation.

Our implementation is also differentiable^7^, thus allowing one to train end-to-end deep learning models with it. Examples of relevant projects are RGN-Networks^8^, or the ongoing open source replication^9^ of AlphaFold2^10^, which makes heavy use of the conversion from internal representation to cartesian coordinates. The differentiability of the code, however, makes it marginally slower because of the data structures needed to accumulate important information for the gradient calculations, and also because the maintenance of this property prevents us from using more efficient libraries like NumPy, that allow compilation of the code to C (e.g. using Cython or Numba) to achieve faster runtimes.

Nevertheless, these features could be adapted for a specific case in which a particular set of properties might be preferred over other ones (e.g. single-thread speed over differentiability, parallelization over single-thread speed, etc.). We leave further optimizations and adaptations to more specific scenarios to the community. In aggregate, we estimate that the current CPU runtimes could be reduced 2x to 3x by adopting a scheme focused on single-thread speed above everything else, which we find to be an acceptable tradeoff given the 400-1200x speedup obtained through parallelism.

## Conclusions

In this work, we have proposed a new, massively parallel scheme for the implementation of the Natural Extension of Reference Frame when applied to polymers, and showcased substantial runtime improvement over previous works. The design principles put in practice allow for easy adoption and usage across the community for different kinds of polymers such as proteins, nucleic acids, glycans or synthetic materials. We hope this accelerated implementation can reduce the times for computational simulations, accelerate the training of machine learning models, and open the window to new advances in polymer structural science.

