## Supplementary material for "MP-NeRF: A Massively Parallel Method for Accelerating Protein Structure Reconstruction from Internal Coordinates": Suplemental Information

### Supplementary Information

The hardware and software used for the experiments in this paper is listed below:

- Laptop:
  - MacBook Pro, intel i5 @ 2.4 GHz and Turbo Boost @ 4.2 GHz. 16 Gb of RAM @ 2133 MHz.
  - MacOS Mojave, Python 3.7.10, Torch 1.8.1
- Desktop: Windows
  - Intel i7-6700K @ 4.00 GHz, 16 Gb of RAM @ 2133 MHz, GPU: Nvidia 1060 GTX 6 Gb
  - Windows 10, Python 3.7.9, Torch 1.7.1

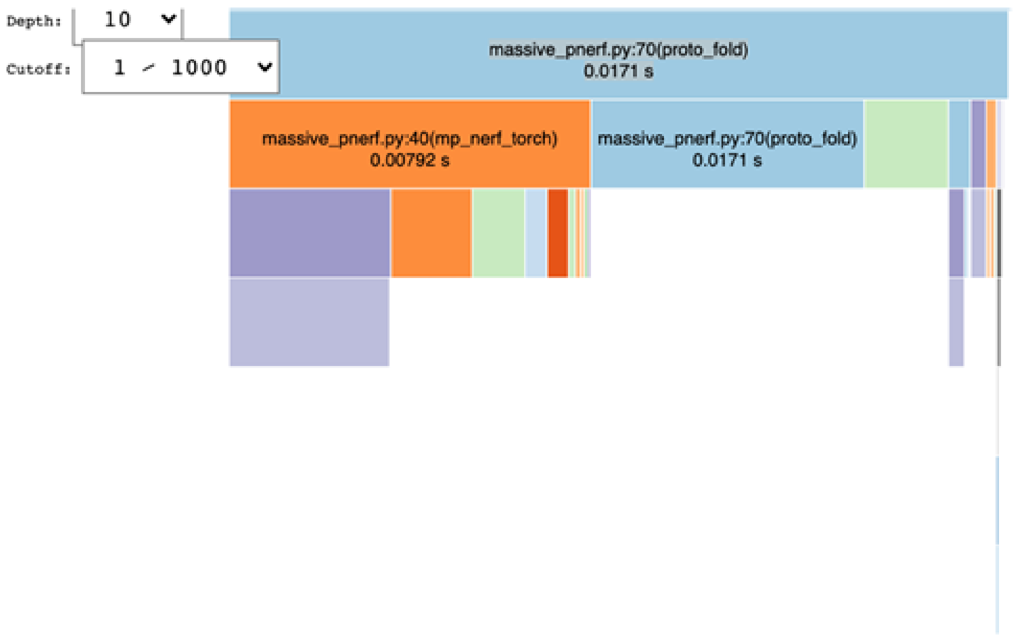


*Figure S1*. Profiling screen capture for the mp-NERF algorithm when folding a protein of 759 AAs. It should be noted that profiling can slow down the execution, the real time being 15.3 ms when no profiling is used, and 17.1 ms when under profiling. Detailed runtimes for the different function calls are disclosed in *table S1*.

| ncalls | tottime | percall | cumtime | percall_1 | filename:lineno(function) |
|---|---|---|---|---|---|
| 1 | 598 | 598 | 171 | 171 | massive_pnerf.py:70(proto_fold) |
| 16 | 4091 | 2557 | 4091 | 2557 | ~:0(<built-in method frobenius_norm>) |
| 773 | 2993 | 0,003872 | 2993 | 0,003872 | ~:0(<built-in method matmul>) |
| 14 | 1775 | 1268 | 792 | 5657 | massive_pnerf.py:40(mp_nerf_torch) |
| 32 | 533 | 0,01666 | 533 | 0,01666 | ~:0(<built-in method cross>) |
| 30 | 495 | 0,00165 | 495 | 0,00165 | ~:0(<built-in method stack>) |
| 14 | 146 | 0,01043 | 146 | 0,01043 | ~:0(<method 'squeeze' of 'torch._C._TensorBase' objects>) |
| 29 | 119 | 0,004103 | 119 | 0,004103 | ~:0(<built-in method cos>) |
| 16 | 117 | 0,007312 | 4243 | 2652 | functional.py:1274(norm) |
| 25 | 103 | 0,000412 | 103 | 0,000412 | ~:0(<method 'unbind' of 'torch._C._TensorBase' objects>) |
| 43 | 1 | 0,002326 | 1 | 0,002326 | ~:0(<built-in method sin>) |
| 14 | 0,00072 | 0,005143 | 0,00072 | 0,005143 | ~:0(<method 'all' of 'torch._C._TensorBase' objects>) |
| 25 | 0,00071 | 0,000284 | 225 | 0,000009 | tensor.py:575(__iter__) |
| 2 | 0,00059 | 0,00295 | 483 | 2415 | massive_pnerf.py:10(get_axis_matrix) |
| 29 | 0,00049 | 0,000169 | 0,00049 | 0,000169 | ~:0(<method 'unsqueeze' of 'torch._C._TensorBase' objects>) |
| 2 | 0,00043 | 0,00215 | 0,00043 | 0,00215 | ~:0(<method 'repeat' of 'torch._C._TensorBase' objects>) |
| 1 | 0,00004 | 0,00004 | 1716 | 1716 | ~:0(<built-in method builtins.exec>) |
| 25 | 0,00003 | 0,000012 | 0,00003 | 0,000012 | ~:0(<built-in method torch._C._get_tracing_state>) |
| 4 | 0,00028 | 0,000007 | 0,00028 | 0,000007 | ~:0(<built-in method tensor>) |
| 6 | 0,00028 | 0,004667 | 0,00028 | 0,004667 | ~:0(<method 'type' of 'torch._C._TensorBase' objects>) |
| 1 | 0,00023 | 0,00023 | 0,00023 | 0,00023 | ~:0(<built-in method zeros>) |
| 1 | 0,00023 | 0,00023 | 1712 | 1712 | <string>:1(<module>) |
| 1 | 0,00019 | 0,00019 | 0,00019 | 0,00019 | ~:0(<built-in method cumsum>) |
| 14 | 0,00018 | 0,001286 | 0,00018 | 0,001286 | ~:0(<method 'item' of 'torch._C._TensorBase' objects>) |
| 11 | 0,00016 | 0,001455 | 0,00019 | 0,001727 | tensor.py:568(__len__) |
| 2 | 0,00015 | 0,000075 | 0,00015 | 0,000075 | ~:0(<built-in method rsub>) |
| 4 | 0,00013 | 0,000325 | 0,00013 | 0,000325 | ~:0(<method 'reshape' of 'torch._C._TensorBase' objects>) |
| 2 | 0,00012 | 0,000006 | 107 | 0,00535 | einops.py:202(apply) |
| 25 | 0,00011 | 0,0000044 | 0,00011 | 0,0000044 | ~:0(<built-in method builtins.iter>) |
| 16 | 0,00011 | 0,0006875 | 0,00002 | 0,000125 | _VF.py:25(__getattr__) |
| 52 | 0,00011 | 0,0002115 | 0,00011 | 0,0002115 | ~:0(<method 'dim' of 'torch._C._TensorBase' objects>) |
| 2 | 0,00001 | 0,000005 | 0,00025 | 0,00125 | tensor.py:525(__rsub__) |
| 2 | 0,00001 | 0,000005 | 12 | 0,00006 | einops.py:327(reduce) |
| 16 | 0,000009 | 0,0005625 | 0,000009 | 0,0005625 | ~:0(<built-in method builtins.getattr>) |
| 34 | 0,000009 | 0,0002647 | 0,000009 | 0,0002647 | ~:0(<built-in method builtins.isinstance>) |
| 1 | 0,000008 | 0,000008 | 0,000008 | 0,000008 | ~:0(<built-in method cat>) |

| | | | | | |
|---|---|---|---|---|---|
| 52 | 0,000008 | 0,0001538 | 0,000008 | 0,0001538 | ~:0(<built-in method torch._C._has_torch_function_unary>) |
| 2 | 0,000006 | 0,000003 | 0,000006 | 0,000003 | ~:0(<built-in method unsqueeze>) |
| 1 | 0,000006 | 0,000006 | 0,000006 | 0,000006 | ~:0(<method 'nonzero' of 'torch._C._TensorBase' objects>) |
| 2 | 0,000006 | 0,000003 | 0,000006 | 0,000003 | ~:0(<method 'permute' of 'torch._C._TensorBase' objects>) |
| 2 | 0,000005 | 0,000025 | 0,000007 | 0,000035 | _backends.py:22(get_backend) |
| 2 | 0,000004 | 0,000002 | 0,000004 | 0,000002 | einops.py:26(_reduce_axes) |
| 2 | 0,000004 | 0,000002 | 0,00059 | 0,00295 | _backends.py:98(add_axes) |
| 2 | 0,000004 | 0,000002 | 0,00047 | 0,00235 | _backends.py:336(tile) |
| 2 | 0,000004 | 0,000002 | 0,000004 | 0,000002 | ~:0(<method 'to' of 'torch._C._TensorBase' objects>) |
| 1 | 0,000004 | 0,000004 | 0,000004 | 0,000004 | ~:0(<method 't' of 'torch._C._TensorBase' objects>) |
| 2 | 0,000003 | 0,000015 | 123 | 0,00615 | einops.py:427(repeat) |
| 2 | 0,000003 | 0,000015 | 0,000009 | 0,000045 | _backends.py:330(transpose) |
| 1 | 0,000003 | 0,000003 | 0,000003 | 0,000003 | ~:0(<method 'view' of 'torch._C._TensorBase' objects>) |
| 6 | 0,000002 | 0,0003333 | 0,000002 | 0,0003333 | ~:0(<method 'items' of 'dict' objects>) |
| 4 | 0,000002 | 0,0000005 | 0,00015 | 0,000375 | _backends.py:83(reshape) |
| 2 | 0,000002 | 0,000001 | 0,000002 | 0,000001 | _backends.py:302(is_appropriate_type) |
| 2 | 0,000002 | 0,000001 | 0,000008 | 0,000004 | _backends.py:339(add_axis) |
| 8 | 0,000001 | 0,0000125 | 0,000001 | 0,0000125 | ~:0(<built-in method builtins.len>) |
| 2 | 0,000001 | 0,0000005 | 0,000001 | 0,0000005 | ~:0(<built-in method builtins.sorted>) |
| 1 | 0,000001 | 0,000001 | 0,000001 | 0,000001 | ~:0(<method 'disable' of '_lsprof.Profiler' objects>) |
| 2 | 0 | 0 | 0 | 0 | ~:0(<built-in method builtins.callable>) |
| 2 | 0 | 0 | 0 | 0 | _backends.py:79(shape) |
| 2 | 0 | 0 | 0 | 0 | ~:0(<built-in method torch._C._has_torch_function_variadic>) |

Table S2. Detailed runtimes for the different function calls in the mp-NERF algorithm. Showing 1 to 59 of 59 entries
